# Classification of ovarian cancer cell lines using transcriptional profiles defines the five major pathological subtypes

**DOI:** 10.1101/2020.07.14.202457

**Authors:** B. M. Barnes, L. Nelson, A. Tighe, R. D. Morgan, J. McGrail, S. S. Taylor

## Abstract

Epithelial ovarian cancer (EOC) is a heterogenous disease consisting of five major pathologically distinct subtypes: High-grade serous ovarian carcinoma (HGSOC), low-grade serous (LGS), endometrioid, clear cell and mucinous carcinoma. Although HGSOC is the most prevalent subtype, representing approximately 75% of cases, a 2013 landmark study from Domcke *et al*., found that many frequently used ovarian cancer cell lines were not genetically representative of HGSOC tissue samples from The Cancer Genome Atlas. Although this work subsequently identified several rarely used cell lines to be highly suitable as HGSOC models, cell line selection for ovarian cancer research does not appear to have altered substantially in recent years. Here, we find that application of non-negative matrix factorisation (NMF) to the transcriptional profiles of 45 commonly used ovarian cancer cell lines exquisitely clusters them into five distinct classes, representative of the five main subtypes of EOC. This methodology was in strong agreement with Domcke *et al*., in identification of cell lines most representative of HGSOC. Furthermore, this robust classification of cell lines, including some previously not annotated or miss-annotated in the literature, now informs selection of the most appropriate models for all five pathological subtypes of ovarian cancer. Furthermore, using machine learning algorithms trained using the classification of the current cell lines, we are able provide a methodology for future classification of novel EOC cell lines.

## Introduction

Ovarian cancer is the most common cause of gynaecological-related cancer death in Europe and North America (Bray et al., 2018). Epithelial ovarian cancer (EOC), which accounts for 80% of all ovarian tumours, is now considered to be a heterogeneous disease consisting of five main histological subtypes characterised by different clinical and molecular features (Lheureux et al., 2019). High-grade serous ovarian carcinoma (HGSOC) is the most prevalent group, accounting for approximately 75% of cases, while the remaining 25% are made up of low-grade serous (LGS), endometrioid, clear cell and mucinous carcinoma (Kurman et al., 2014). Endometrioid and mucinous carcinoma are further sub-classified into well, moderately and poorly differentiated tumours (grade 1 to 3, respectively) (Kurman et al., 2014). Diagnosis of each subtype of EOC involves histological examination in combination with immunohistochemistry analysis, which is considered gold standard (Kurman et al., 2014).

Expansion of next generation sequencing has enabled closer inspection of the unique genomes of each subtype of EOC. HGSOC are characterised by near-ubiquitous *TP53* mutation and genome-wide copy-number variation (CNV), with germline or somatic *BRCA1/2* variants present in ∼ 20% of cases (Bell et al., 2011; Ciriello et al., 2013; Huang et al., 2018). LGS less frequently shows *TP53* mutation, and instead variants in the MAPK signalling pathway are observed (e.g. *KRAS, NRAS, BRAF*) (Etemadmoghadam et al., 2017; Fernandez et al., 2019; Jones et al., 2012). Clear cell carcinomas and well-differentiated (i.e., grade 1) endometrioid carcinomas are commonly associated with endometriosis and *ARID-1A* variants (Jones et al., 2010; Wiegand et al., 2010). Finally, mucinous ovarian carcinoma is associated with *KRAS* variants and *ERBB2* amplifications (Cheasley et al., 2019).

Cancer cell lines are often used as model systems to study cancer; however, most were established many years ago and have either genetically drifted from the original patient cells and/or lack sufficient clinical data to allow robust tumour type classification. For example, much of ovarian cancer research has been based on the SKOV-3 cell line, however an in-depth analysis of copy-number changes, mutations and microarray-based mRNA expression profiles revealed that this cell line and others are actually atypical, bearing few hallmarks of the most common type of ovarian cancer, HGSOC, as defined by comparison with patient samples from The Cancer Genome Atlas (Bell et al., 2011; Domcke et al., 2013). Indeed, this analysis by Domcke *et al*. represented a landmark in the field, identifying a number of Cancer Cell Line Encyclopaedia (CCLE) cell lines that better reflect the genomic and mRNA expression landscapes of HGSOC.

This raises a key question: without directly associated clinical and/or histopathological annotation, how does one determine which of the subtypes any given cell line or patient biopsy reflects? Here we set out to address this question by asking whether it is possible to distinguish EOC subtypes based on molecular fingerprints, in particular one derived from RNA-sequencing (RNAseq). While the utility of RNAseq as a tool for developing prognostic biomarkers is still in its infancy, the technique is tried and tested, has the potential to provide a wealth of information by interrogating the expression levels of tens of thousands of genes and is gradually becoming more accessible and less costly. The challenge is in the distilling of robust signatures that correlate with specific phenotypes from these complex datasets.

One approach to reducing the complexity of RNAseq data is non-negative matrix factorisation (NMF), which has been utilised to reduce the dimensionality of transcriptional profiles from thousands of genes to a subset of important metagenes, concurrently providing meaningful class discovery (Brunet et al., 2004). Here, we apply NMF to the gene expression profiles of 45 EOC cell lines sequenced as part of the CCLE. We demonstrate the decomposition of this panel of EOC cell lines into five robust clusters that recapitulate the characteristics of the different pathological histotypes. In turn, this allows reclassification of several cell lines that were previously not annotated or possibly miss-annotated. Our results align well with the analysis by Domcke et al., which was based on CCLE’s earlier microarray gene expression dataset. Our analysis further facilitates selection of cell lines appropriate for research of HGSOC, and in addition identifies cell lines representing the other four EOC subtypes. We also provide a methodology for future classification of novel cell lines using a K-nearest neighbour (KNN) classifier trained on the CCLE cell lines.

## Results and Discussion

### Most frequently utilised CCLE lines are unlikely to be representative of HGSOC

The analyses by Domcke *et al*. represents an important milestone in the field, ranking 47 ovarian cancer cell lines according to their genetic and gene expression resemblance to HGSOC. In the intervening seven years, additional data has become available, in particular RNAseq data. We therefore set out to revisit this issue. Our aim was to determine whether the next generation of gene expression profiling clusters EOC cell lines into the different histotypes by NMF, and evaluate the ability of common machine learning algorithms, KNN, random forest and support vector machine (SVM), trained to identify the NMF-assigned class.

Firstly, we performed an extensive literature search to collate all annotations related to the 47 CCLE cell lines with site of origin indicated to be the ovary (with available RNAseq data). This identified 44 cell lines of EOC origin, eliminating 3 representing the non-epithelial Brenner and granulosa tumour types, and an engineered/immortalised cell line. Information gathered included reported histotype, specimen site, pre-biopsy treatment, the HGSOC likelihood score (as determined by Domcke *et al*.) and any other relevant information, for example, age and clinical course (Table S1). We also determined cell line usage in research by PubMed search (see Table S2 for search terms, including aliases for each cell line). Interestingly cell line selection has not substantially altered in recent years, despite publication of Domcke’s landmark study in 2013. Seven cell lines (ranked by most highly used: SKOV-3, A2780, OVCAR-3, IGROV-1, CAOV-3, 59M and OVCAR-8) collectively constitute almost 90% of the total PubMed citations (Fig. 1). Of these 7, only three received a ‘HGSOC-likely’ score in the analysis by Domcke *et al*. (OVCAR-3, CAOV-3 and 59M). Strikingly, seven cell lines scoring highly as ‘HGSOC-likely’, KURAMOCHI, OVSAHO, SNU-119, COV362, OVCAR-4, COV318 and JHOS-4, only constitute 1.07% of PubMed usages of the 44 EOC cell lines included in the CCLE. Furthermore, as late as 2019, SKOV-3 and A2780 remain the first and second most highly studied cell lines in ovarian cancer research, respectively, despite their purported unsuitability as HGSOC cell line models.

**Figure 1.**
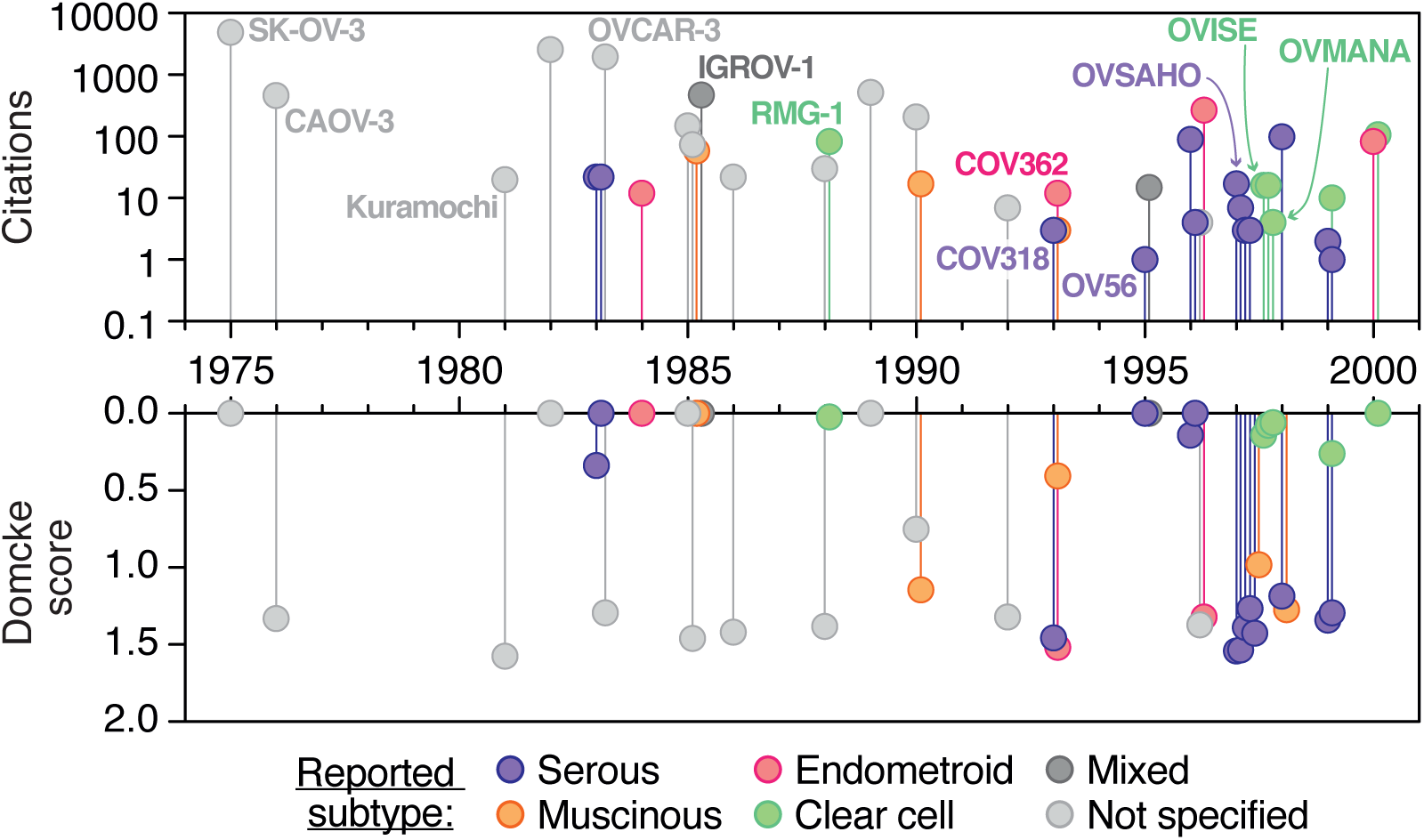
Cell line usage based on PubMed citations. Top, total number of PubMed usages of each of the epithelial ovarian cancer cell lines for which RNAseq data is available within the CCLE. Bottom, HGSOC-likelihood scores as determined by Domcke et al. analysis of ovarian cancer cell lines correlated with The Cancer Genome Atlas HGSOC patient samples. Cell lines are separated along the x-axis based on the year of their first usage. Cell lines are coloured by the subtype of epithelial ovarian cancer reported in their primary literature source. Green, clear cell; red, endometrioid; orange, mucinous; purple, serous; dark grey, mixed; light grey, not specified (NS).

### Cancer cell lines cluster into classes representative of the five EOC histotypes

Next we obtained from the European Nucleotide Archive the raw RNAseq files for the 44 EOC cell lines analysed by the recent CCLE project (Ghandi et al., 2019) and mapped reads to the GRCh38 human genome assembly with gene annotations from Gencode v32. The most important parameter to estimate in any clustering method is the optimum number of clusters (k) for the data. The consensus matrix methodology by Monti et al. (2003) is frequently used in the evaluation of clustering, where the entries of the consensus map are coloured from 0 to 1, reflecting the probability of clustering of two samples together across multiple runs of NMF (see Fig. S1 for consensus maps of all NMF models from k of 2 to 7).

Many quality metrics have been proposed to assess the optimum value of k (Fig. 2A): briefly, Brunet et al. (2004) proposed the cophenetic correlation coefficient, Kim and Park (2007) proposed the dispersion coefficient, Rousseeuw *et al*. (1987) proposed the silhouette width. In each instance, the value of k that results in maximum of the coefficient is chosen as optimum. Additionally, Hutchins et al (2008) utilised the variation of the residual sums of squares (RSS) between the original data and estimated data (not shown). The value of k at which the plot of RSS for each value of k shows an inflection point can be chosen as the optimum. Plotting these metrics for 2 to 10 clusters revealed that both two and five clusters fitted the dataset well (Fig. 1A). However, at a factorisation rank of two, no biologically interpretable clustering was apparent, with cell lines reported as individual subtypes split across the two clusters (Fig. S1a). We backwards annotated each cell line with the cluster assignment from the NMF run using 5 clusters, and performed consensus clustering on the result of the NMF run using just two clusters (not shown). There was no readily observable stratification of the five clusters, or combination thereof, with each of the five clusters split across the two clusters. We inferred, therefore, that there were no nested structures present within the data as k was increased from 2 to 5, as was observed previously in the classification of leukaemia samples using NMF (Brunet et al., 2004). Brunet *et al*. found that at a factorisation rank of 2, ALL and AML samples clustered separately. As the factorisation rank was increased from 2 to 3, the ALL cluster divided into the T-cell and B-cell distinctions. Thus, NMF has been reported to reveal hierarchical structure when it exists, without forcing such structure on the data (as other clustering models may), highlighting the strengths of NMF over other methods (Brunet et al., 2004).

**Figure 2.**
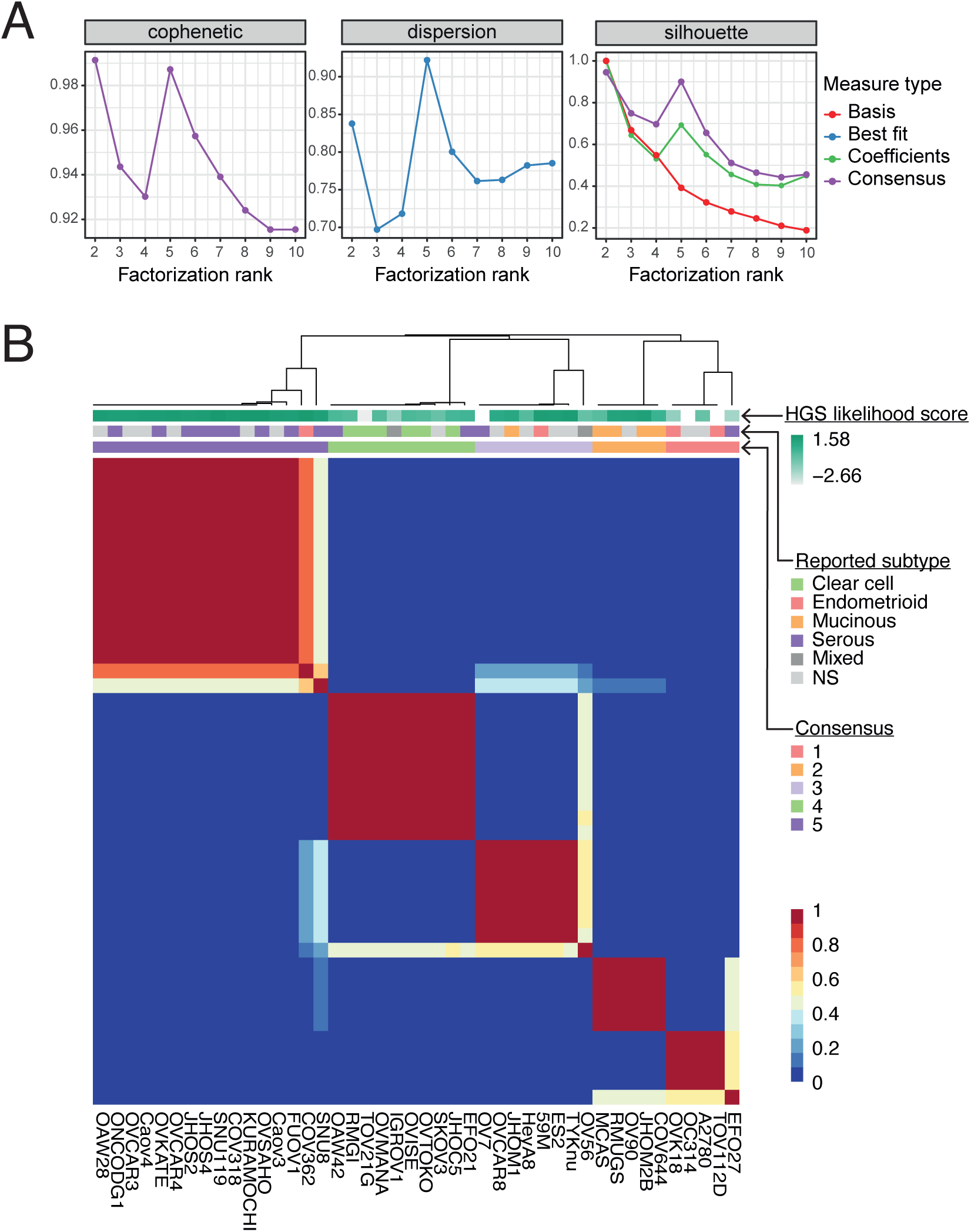
Ovarian cancer cell lines can be divided into five clusters that recapitulate the histological subtypes based on transcriptional profiles. (**A**) Selected quality metrics describing the performance of non-negative matrix factorisation for 2 to 10 clusters. From left, the cophenetic correlation, dispersion and silhouette coefficients. Colours indicate the type of measure plotted. (**B**) Consensus map showing cell line clustering for 200 iterative runs of NMF using 5 clusters. The blocks of the consensus map are coloured by the probability of two samples clustering together, where red, 1; white, 0.5 and blue, 0. The annotation track atop the heatmap indicates (top) the HGSOC-likelihood score of a cell line determined by Domcke et al. Where darker shades represent a higher score. The pure white blocks indicate the cell line was not included in this analysis. Middle track, the ovarian cancer subtype provided in the cell line’s original literature source where green, clear cell; red, endometrioid; orange, mucinous; purple, serous; dark grey, mixed; light grey, not specified (NS). Bottom track, the consensus cluster assignment across 200 NMF runs where dark purple, cluster 1; green, cluster 2; light purple, cluster 3; orange, cluster 4 and red, cluster 5.

In the CCLE EOC dataset, NMF together with consensus clustering gave strong evidence for a five-class split with clear block diagonal patterns and correspondingly high-quality metrics, with k=5 cophenetic and silhouette width scores second only to k=2 (Fig. 2A). However, the dispersion score was highest for k=5 (Fig. 2A), and the RSS curve shows an inflection point at k=5 (not shown), tying k=2 and k=5 as the optimum. We then examined the subtype assigned by the primary literature source for each cell line (where available; Table S1). Interestingly, this showed a clear overrepresentation of cell lines from a given subtype contained within each cluster, suggesting that the clusters identified by NMF are representative of the major EOC subtypes of ovarian cancer (Fig. 1B).

#### High grade serous ovarian carcinoma

We begin our discussion of the five clusters with the top left of the consensus map (Fig. 2B; dark purple). Of the cell lines in this cluster, 8 of 16 were assigned ‘serous’ in their primary literature annotation. Of the remaining 8 cell lines, 1 was reported as endometrioid (COV362) and the subtype of the remaining 7 was not specified in the literature. Given the putative identification of this cluster as representing HGSOC-derived cell lines, we wanted to align our results with the likelihood scores of these cell lines determined in the analysis by Domcke *et al*. (Fig. 1B; blue/green graduated track). In fact, all 16 cell lines that fall within this cluster were within the top 20 scoring cell lines in the previous analysis, providing remarkable confirmation of the methodology used here and by Domcke *et al*. for annotating cell lines as representative of HGSOC. Of the cell lines not placed into the HGSOC cluster, but ranked in the top 20 of Domcke *et al*., TYK-nu and 59M were designated ‘likely HGS’ and JHOM-2B and ES2 ‘possibly HGSOC’. We discuss these cell lines in the context of their assigned cluster in the relevant sections below. Therefore, clustering, confirmed several cell lines without specified subtype in their primary literature source, to represent good models of HGSOC, including KURAMOCHI, OVCAR-4, Caov-4, OAW28, Caov-3, ONCO-DG-1, and OVCAR-3. The cell lines OVSAHO, SNU-119, COV318, JHOS-4, JHOS-2, OVKATE, FU-OV-1 and SNU-8 retained their literature classification as ‘HGSOC’ in our analysis.

COV362 was initially annotated as endometrioid in the literature, however here we find it clusters with the cell lines representing HGSOC. This line has a *TP53* mutation and a *BRCA2* mutation, lesions characteristic of HGSOC, supporting the placement of COV362 as HGSOC. However, it should be noted that SNU8 and, to a lesser extent, COV362, show disparate clustering across 200 runs of NMF with random initialisation points. COV362 also clustered 25% of the time into cluster 3 (low grade serous), suggesting that it may share some characteristics of these cell lines. Importantly, it does not cluster in any of the NMF runs with other cell lines reported as endometrioid, further suggesting that this designation may be incorrect. SNU8 also clustered in approx. 42% of NMF runs with cluster 3 (low grade serous) and in 14% with cluster 4 (mucinous)

#### Clear cell

In the next cluster (second from the left; green), there is an enrichment of cell lines which were defined as clear cell in their primary literature source. In fact, of the 10 cells lines, 6 were annotated as clear cell in the original publication, 2 were annotated as serous, 1 mixed and 1 was not specified. No cell lines annotated primarily as clear cell in the literature fell into any other cluster. The two samples previously annotated as serous were EFO21 and OAW42. Indeed, both of these cell lines received relatively low HGSOC likelihood scores in the analysis by Domcke *et al*., suggesting they are poor HGSOC models. Unlike almost all HGSOC, OAW42 has wild-type *TP53*. However, it does harbour two separate frameshift mutations within *ARID1A*, supporting its designation here as clear cell (Wiegand et al., 2010). Although EFO21 has mutated *TP53*, and no ARID1A mutation, these cells have amplification of PIK3CA, showing resultant mRNA expression levels within the 93rd percentile of CCLE cell lines. The most common mutations identified by sequencing of a 46 gene panel using pure clear cell samples included mutations in *PIK3CA* (50.0%; 52 of 104 cases tested), *TP53* (18.1%; 19/105), and *KRAS* (12.4%; 13/105) (Friedlander et al., 2016). Our analysis therefore also supports EFO21 classification as a clear cell line.

The most heavily used ovarian cancer cell line, SKOV-3, also falls within this cluster. Despite its extensive use, the primary literature source does not designate SKOV-3 to any particular subtype. Interestingly, SKOV-3 may actually be one of the most typical examples of clear cell as they harbour aberrations of three of the most commonly mutated proteins in clear cell ovarian cancer: *PIK3CA, ARID1A* and *TP53*. Therefore, designation here as clear cell is most likely an accurate representation of this cell line.

#### Low grade serous

In our analysis TYK-nu and 59M cluster together in cluster 3, which we believe to represent LGS. The CCLE/broad institute report TYK-nu as having a *TP53* mutation, which molecular studies of LGS suggest are less common in this subtype (8% in LGS versus 96% in HGSOC) (Bell et al., 2011; Singer et al., 2005). However, LGS is also characterized by activation of the mitogen-activated protein kinase (MAPK) pathway. Mutations affecting this pathway are seen in *KRAS, NRAS* and *BRAF* genes, in addition to multiple alterations affecting other genes related to this pathway (Etemadmoghadam et al., 2017; Fernandez et al., 2019; Jones et al., 2012). In addition, copy number alterations and mutations affecting 61 MAPK-related genes were recently identified in 14 LGS cell lines (Fernandez et al., 2019). In this vein, TYK-nu have two mutations within *NRAS*, a member of the RAS/RAF pathway not included within Domcke’s scoring schema. Furthermore, TYK-nu is derived from a 38-year-old patient in line with reports that LGS affects women at a younger age than HGSOC, with a median age at diagnosis for LGS of between 43 and 47 years (Gershenson, 2016; Gershenson et al., 2015). 59M, while also harbouring a *TP53* mutation, has three mutations in proteins in the MAPK pathway (Ghandi et al., 2019), and is therefore characteristic of LGS (previously annotated as endometroid). (Wilson et al., 1996)

The group of Coscia *et al*. used a proteomic signature to stratify putative HGSOC cell lines into three distinct groups (Coscia et al., 2016). Although the majority of cell lines with a high genetic fidelity to HGSOC were classified as group I and bore a more epithelial proteome, the two cell lines that clustered in group III with a more mesenchymal proteome were 59M and TYK-nu. While there was a striking concordance between the proteomic signature of group I cell lines and HGSOC patient samples, as well as cultured fallopian tube epithelial cells, group III cell lines resembled the signature of immortalized ovarian surface epithelial cells. Although the authors suggest that heterogeneity exists in the proteome of HGSOC based on disparate sites of origin (Coscia et al., 2016), it could be argued that these differences actually represent the differences between HGSOC and LGS-derived cell lines.

Collectively, this suggests TYK-nu and 59M form part of a cluster of 8 LGS cell lines (Fig. 1B; light purple). As LGS represents a fairly recent descriptor, it is difficult to infer this annotation from primary literature annotations of cells lines. Here we identify 4 cell lines, TYK-nu, HeyA8, ES2, and OVCAR8, which were previously unspecified in the literature, to be representative of LGS. In addition, JHOM-1 also clusters here, which was initially annotated as mucinous in its primary literature source.

#### Mucinous

Of five cancer cell lines annotated in their primary reference as mucinous, four of them fall into cluster number 4. These are MCAS, RMUG-S, COV644 and JHOM-2B. Of the cell lines determined to be in the top 20 of HGSOC likely cell lines by Domcke *et al*., JHOM-2B is reported in the literature as mucinous and our NMF also clusters it with the majority of other mucinous cell lines, suggesting its original classification is correct. In fact, Domcke et al. ranked JHOM-2B as 19th, close to the threshold for designation as ‘possibly HGS’. Indeed, this cell line does harbour a *TP53* mutation, which may disproportionately influence its standing in the analysis by Domcke *et al*. However, while *TP53* mutations are almost ubiquitous in HGSOC ovarian cancer, around 16% of mucinous tumours show mutated *TP53* (Schuijer & Berns, 2003). The fifth cell line reported as mucinous in its original publication is JHOM-1, falls into the cluster we tentatively class as LGS (discussed previously).

The cell line OV-90 also clusters with the mucinous cell line, which originally was not designated a subtype in the original articles. In support of its mucinous designation, it harbours *ERB2* amplification and *BRAF* mutation which have been demonstrated in mucinous ovarian cancer (Cheasley et al., 2019; Friedlander et al., 2015).

#### Endometrioid

Finally, the fifth cluster, designated endometroid, is constituted of two cell lines that were annotated as such in their primary reference, namely TOV112D and OVK18. Two other cell lines annotated as endometroid in their primary reference fall into cluster 3 (which we tentatively label as the LGS cluster; 59M) and cluster 1 (HGSOC cluster; COV362), and their suitability to fit these clusters has been discussed previously. Two further cell lines that cluster as endometroid here, A2780 and OC314, were not assigned a subtype in their primary literature source and are therefore newly annotated as potential models of endometroid ovarian cancer.

Lastly, EFO-27 also clusters within the endometroid cluster. Although this cell line was originally classified as serous in the literature, it received a poor HGS-likelihood score in the work by Domcke *et al*., giving initial evidence of its unsuitability as a HGSOC model cell line. EFO27 cells harbour a missense mutation in *PPP2R1A*, which has previously been found to be mutated in 12.2% (5/41) of endometrioid ovarian cancers, but not in 50 high-grade and 12 low-grade serous carcinomas (McConechy et al., 2011). More recent genetic screens of endometrioid ovarian cancer identified similar driver mutations to endometrial carcinoma, including *PTEN, CTNNB1, PIK3CA, ARID1A, TP53, KMT2D, KMT2B* and *PIK3R1* (Pierson et al., 2020). Indeed, with the exception of *CTNNB1*, EFO-27 have mutations in all these genes (Ghandi et al., 2019). Therefore, the genetic similarities between EFO-27 and endometrioid ovarian cancer support it representing a better model of this type of ovarian cancer, than of HGSOC. However, it should be noted that this cell line has a poor silhouette score in our consensus map (Fig. 2B), clustering with other endometrioid cell lines 58% of the time, and with cluster 4 (the mucinous cluster) in the other NMF runs. Of the genetic lesions associated with mucinous ovarian cancer (Friedlander et al., 2015), EFO-27 harbours *PTEN* and *PIK3CA*. This cell line does not harbour *KRAS* mutation or *ERBB2* amplification, however, which have been shown to be mutated in mucinous ovarian cancer (Cheasley et al., 2019).

### Evaluating machine learning algorithms to classify ovarian cancer subtypes

We next sought to determine whether the NMF class given to each cell line could be used to train a machine learning model to predict the subtype of a ‘hold-out’ set. Genes whose expression levels were characteristic for each cluster were extracted with each cluster containing between 23 and 82 such metagenes. The largest number of metagenes was associated with the putative HGSOC cluster (82), followed by, endometrioid (40), LGS (35), mucinous (28) and clear cell (23) (Fig. 3A). We next evaluated the classification potential of several common machine learning algorithms: KNN, random forests and SVM. The 45 cell lines were randomly partitioned into four groups, such that each group had an even representation of cell lines from each subtype. Then, each model was trained to each successive set of 3 groups, and model performance tested on the omitted group. This meant that each sample had an opportunity to be both trained and tested on. The per-subtype specificity and sensitivity metrics were compared across KNN, random forest and SVM algorithms (Fig. 3B). As can be seen, all models predicted the HGSOC subtype well, achieving balanced accuracy scores of 1 (KNN), 0.935275 (RF) and 0.984375 (SVM) for this class. This presumably reflects the larger number of samples labelled HGSOC and the number of metagenes present to predict this subtype versus the others. Therefore, additional samples representative of non-HGSOC ovarian cancer would greatly aid the training of a classifier. This is especially true in the case of endometrioid ovarian cancer cell lines, which was represented by only 4 of the 44 cell lines analysed in this study. Nevertheless, the overall kappa values achieved for each model was 0.918 (KNN), 0.78905 (RF) and 0.878 (SVM). This suggests that NMF coupled with KNN may be a powerful tool for ovarian cancer cell line subtype classification.

**Figure 3.**
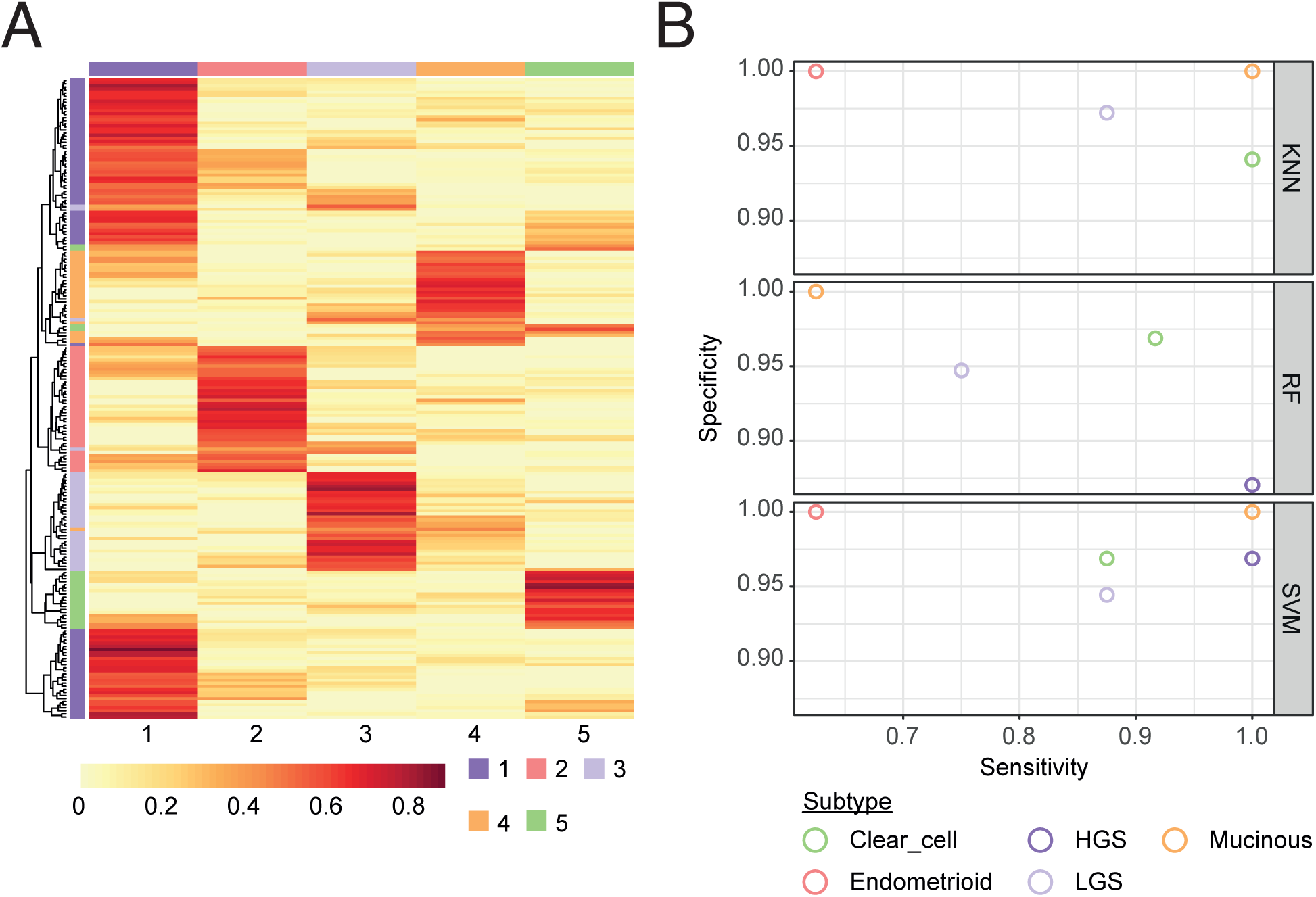
A k-nearest neighbour classifies accurately predicts subtype of ovarian cancer cell lines. (**A**) Metagenes for which high expression is informative of each cluster were extracted using gene scoring scheme as per Kim and Park (2005). Colours represent the strength of the association between that gene and the cluster, where red, 1 and white, 0. The track above the heatmap indicates cluster number, as per Fig. 2, where dark purple, cluster 1; green, cluster 2; light purple, cluster 3; orange, cluster 4 and red, cluster 5. (**B**) Evaluation of three machine learning algorithms for ovarian cancer cell line subtype classification, k-nearest neighbour (KNN), random forest (RF) and support vector machine (SVM). Cell lines were designated the subtype indicated by NMF clustering, and partitioned into 4 subsets. Three subsets were used to train each of the machine learning algorithms, with the fourth set held out as a test set. The four subsets were rotated such that each sample had the opportunity to be trained and tested upon. The average per-class sensitivity and specificity scores across the four tested sets is shown where dark purple, HGSOC; green, clear cell; light purple, LGS; orange, mucinous and red, endometrioid.

## Conclusion

The EOC subtype from which commonly used ovarian cancer cell lines were derived has remained a controversial topic for many years (Anglesio et al., 2013; Beaufort et al., 2014; Coscia et al., 2016; Domcke et al., 2013). We sought to determine whether recently released RNAseq data from the CCLE could shed light on this subject. Previous studies have sought to define an immunohistochemical, genetic or combinatorial panel, and determine the suitability of cells to fit this mould. Here we have not imposed any prior knowledge or structure onto the data, instead opting to use NMF, a clustering algorithm that has been used not only in gene expression studies, but other pattern-recognition problems such as facial recognition and deciphering the meaning of words (Brunet et al., 2004; Lee & Seung, 1999). Our NMF clustering allowed cell lines to cluster with others that they most closely resembled at the transcriptional level, revealing novel subtype classifications for some cell lines. Inclusion of additional cell lines would improve the predictive utility of our machine-learning based classifier, especially subtypes that are underrepresented in the CCLE dataset, namely endometrioid and mucinous. Future work, therefore, could relate to the integration of multiple different sources of transcriptional profiles. Additionally, datasets containing patient-derived cell lines could be utilised to further evaluate the performance of any classifier, including the recently published living ovarian biobank and others (Fernandez et al., 2019; Nelson et al., 2020).

## Materials and Methods

### Literature search

We performed an extensive PubMed literature search to determine the usage of CCLE ovarian cancer cell lines. The list of search queries used is supplied in table S2, demonstrating the different aliases used for the different cell lines. It should be noted that these search queries only count the number of articles where the cell line name was specified in the title and/or abstract, therefore missing some articles that only specify within article the cell lines used. This will be especially true for larger studies that utilize many of these cell lines where it is not possible to list them in an abstract.

### RNAseq data

Forty five cell lines representative of the major ovarian cancer subtypes analysed by RNA-sequencing as part of the Cancer Cell Line Encyclopedia (CCLE) project (Ghandi et al., 2019) were identified (table S1). Raw sequence files in FASTQ format were obtained from the European Nucleotide Archive (ENA; http://www.ebi.ac.uk/ena/). STAR (v2.7.2a) (Dobin et al., 2013) was used to map reads to the GRCh38 human genome assembly with gene annotations from Gencode v32. The number of reads per gene were counted using --quantMode GeneCounts within the STAR command.

### Non-negative matrix factorisation

Data analyses in R was performed using v3.6.2 and in Bioconductor v3.10. The DESeq2 (v1.26.0) (Love et al., 2014) package was used to apply a variance stabilizing transformation to the assembled read count matrix. Transcripts with a median absolute deviation ≥1.5 were selected, and this list of 6,796 genes was used as input for clustering analysis using the NMF package (Gaujoux & Seoighe, 2010). To estimate the factorisation rank (k), NMF was performed for a k of 2 to 10 using 50 random initiations. Quality measures were computed for each factorisation rank, including the cophenetic coefficients, silhouette and RSS. Inspection of the computed quality metrics revealed 5 clusters fitted the data. Next, 200 iterative runs of NMF were performed from a fixed random initial condition with a k value of 5. Using annotations given in the primary literature source for each cell line (table S1), we inferred the likely ovarian cancer histotype of each cluster. Gene scoring schema was applied to extract genes characteristic of the five identified clusters (Kim & Park, 2007). Metagene lists were combined, and this was used as input for machine learning algorithms.

### Machine Learning Algorithms for Classification

A plethora of classification algorithms have become available. Here, we explore the utility of three common classification algorithms: KNN, RF and SVM. We used the R package caret (v6.0-86) for model training and evaluation. The specific modules used were base::knn, randomForest (v4.6-14) and kernlab (v0.9-29), respectively. The cell lines with their subtype classifications outputted from our NMF analysis were partitioned into 4 random subsets, such that each set contained approximately equal proportions of each subtype. Models were trained using each combination of partitions, leaving one group out for testing of model performance in each instance. Metrics compared between models were the per-class (ability to predict each subtype, e.g. HGSOC, LGS etc.) sensitivity, specificity and balanced accuracy calculations. Overall model performance was compared using Cohen’s kappa, which compares observed accuracy with the expected accuracy (subtypes predicted by a random classifier).

#### K-nearest neighbours

K-nearest neighbours is a non-parametric method proposed by Thomas Cover used for classification. A cell line within the held-out test set is classified by majority vote of its k-neighbours from the training set (although no explicit training step is required). K is typically a small positive integer, and usually of an odd number to avoid ‘tied’ decisions. A large k reduces the impact of variance caused by random error. However, this may miss the small but important patterns within the data (Zhang, 2016).

#### Random Forrest

Random forest is a learning method for classification, regression and other tasks. The forest is built from the construction of many different decision trees at training time. The power of the algorithm stems from the low-correlation between decision trees, which may cancel out the individual errors of any one tree. Each tree decides the subtype of a test-set cell line and the majority vote becomes the model’s prediction. While some trees may be wrong, many other trees will be right, so as a group the trees are able to provide a more powerful prediction.

#### SVM

Support vector machine is a supervised machine learning algorithm that can be employed for both classification and regression purposes. SVM works by finding the decision boundary (the “hyperplane”) that separates the classes of the supplied data, in our case the different subtypes of EOC. During training, the margins of the hyperplanes are maximised, while the cell lines remain on the correct side of the subtype boundaries. Intuitively, when the subtype of the test is predicted, we can be more confident that the prediction is correct if the cell lines lies further from the boundaries. Likewise, doubt is cast on the prediction of a cell line that sits close to the boundaries.

### Genetic background and copy number variation of CCLE cell lines

The genetic background of the CCLE cell lines is extensively referred to throughout this manuscript. We direct the reader to the mutation and copy number variation datasets generated by this project. The datasets were originally presented in Ghandi et al (2019) and recommend the use of the cBioPortal for Cancer Genomics (https://www.cbioportal.org/) that enables interactive exploration of multidimensional cancer genomics data sets (Cerami et al., 2012; Gao et al., 2013).

## Acknowledgments

We thank the members of the Taylor lab for advice and comments on the manuscript. The research was funded by a Cancer Research UK Programme Grant to S.S.T (C1422/A19842) and the Cancer Research UK Centre Award (C5759/A25254).

## Author contributions

Methodology, Investigation, Validation and Formal Analysis, B.B., L.N., A.T. and R.D.M.; Conceptualisation, B.B. and R.D.M; Writing, B.B., R.D.M., J.M. and S.S.T.; Funding and Supervision S.S.T.

## Figure legends

**Figure S1.**
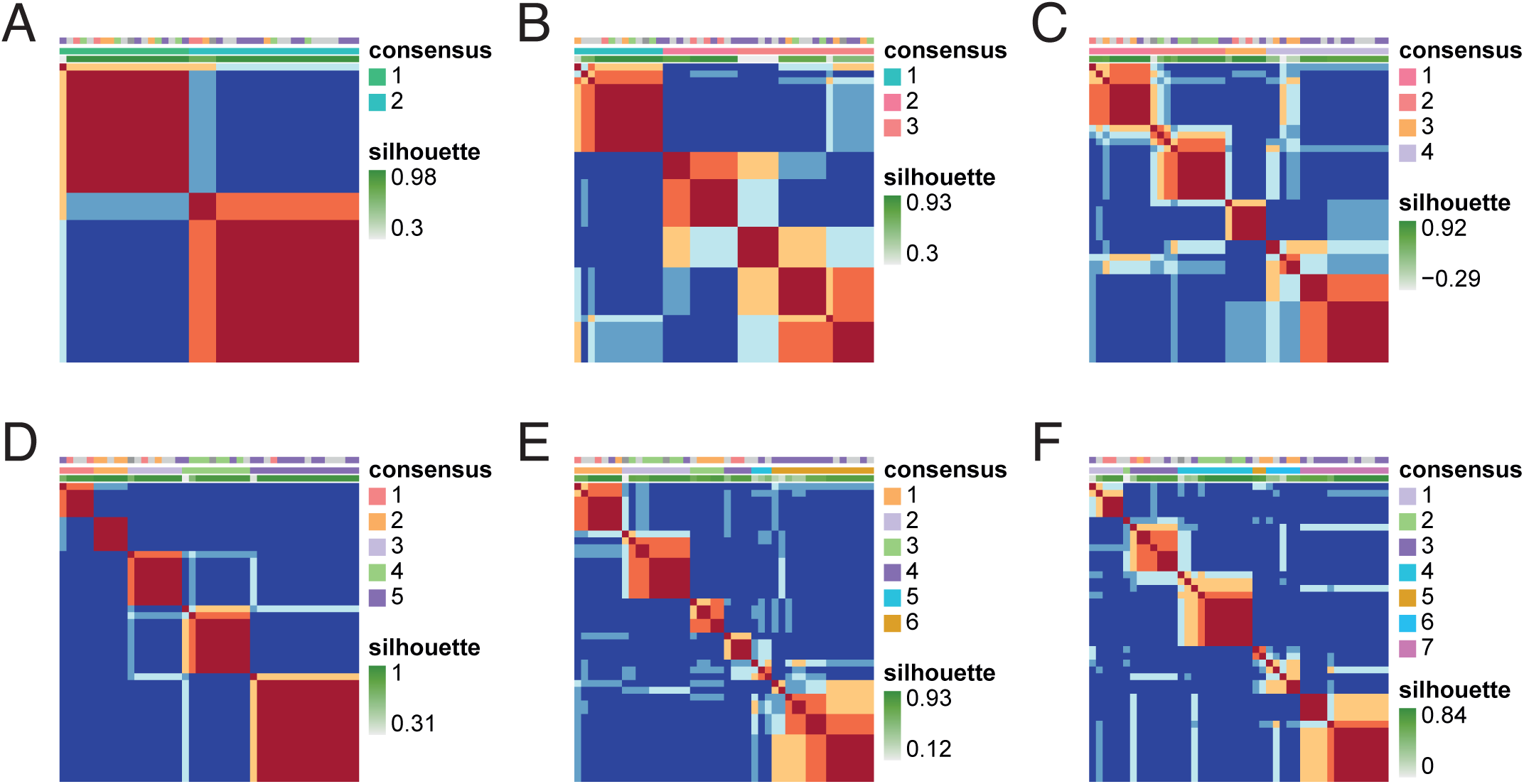
consensus cluster maps for NMF at different values of k. (**A-F**) consensus cluster maps (in order of increasing k) from 2 to 7 clusters. The blocks of the consensus map are coloured by the probability of two samples clustering together, where red, 1; white, 0.5 and blue, 0. The annotation tracks atop the heatmap indicate the ovarian cancer subtype provided in the cell line’s original literature source where green, clear cell; red, endometrioid; orange, mucinous; purple, serous; dark grey, mixed; light grey, not specified (NS). Middle track, the consensus cluster assignment across 50 NMF runs. The cluster numbers and the colours assigned are shown in the legends to the right of each of the heatmaps. Bottom track, silhouette width for each sample pair where dark green indicates a silhouette width of 1 (perfect clustering).

